# Evolution shapes metabolic function and niche-specific antimicrobial targets in pathobionts

**DOI:** 10.1101/2022.11.10.515998

**Authors:** Emma M. Glass, Lillian R. Dillard, Glynis L. Kolling, Andrew S. Warren, Jason A. Papin

## Abstract

Treatment of infections with traditional antimicrobials has become difficult due to the growing antimicrobial resistance crisis, necessitating the development of innovative approaches for deeply understanding pathogen function. Here, we generated a collection of genome-scale metabolic network reconstructions to gain insight into evolutionary drivers of metabolic function. We determined physiological location is a major driver of evolution of metabolic function. We observed that stomach-associated pathobionts had the most unique metabolic phenotypes and identified three essential genes unique to stomach pathobionts across diverse phylogenetic relationships. We demonstrate that inhibition of one such gene, *thyX*, inhibited growth of stomach-specific pathobionts exclusively, indicating possible physiological niche-specific targeting. This pioneering approach is the first step to using unique metabolic signatures to inform targeted antimicrobial therapies.

**One sentence summary:** A data-driven approach to drug target discovery through metabolic signatures of diverse pathogens conserved across body-sites.

## Introduction

Bacterial pathogens pose a major risk to human health, and are responsible for 16% of all global deaths, including 44% of deaths in low-resource countries(*1*). Currently, there are over 500 known human-associated bacterial pathobionts(*2*) -- a group of taxonomically diverse microorganisms that are characterized by their capacity to exhibit pathogenicity(*3*). Treatment of certain bacterial pathobiont infections with traditional antimicrobials has become increasingly difficult in recent years due to growing resistance (*4*). It is therefore necessary to gain a deeper understanding of the evolutionary principles governing pathobiont metabolism to uncover cellular pathways that could be newly exploited for targeted antimicrobial therapies.

Many well-known pathobionts have been deeply characterized experimentally and computationally (*5–7*), but metabolic phenotypes have not been described across pathobiont genera. It is necessary to capture systematically the complex variation in metabolic phenotypes across pathobionts to begin to understand the evolution of unique metabolic function. Understanding this variation in phenotypes could allow for identification of unique metabolic signatures in individual or select groups of pathobionts, such as those inhabiting the same physiological niche. Identifying niche-specific metabolic signatures could allow us to consider antimicrobial drug development from a different perspective; targeting cellular processes that are conserved amongst pathobionts inhabiting a specific body site could reduce the need for broad-spectrum antibiotics, ultimately leading to a reduction in the emergence of antimicrobial resistance.

To capture the range of metabolic phenotypes across pathobionts, we need to leverage a high-throughput, automated, *in silico* pipeline. Specifically, we can use genome-scale metabolic network reconstructions (GENREs) to capture functional metabolism in individual pathobionts, at strain-specific resolution (*5*, *8*). Once assembled, GENREs can be used to probe an organism’s genotype-phenotype relationship through constraint-based modeling and analysis (COBRA)(*9*). *In silico* modeling of bacterial metabolism through GENREs has proven effective at defining functional metabolism in individual pathogens (*5*, *8*, *10*, *11*).

Here, we identify evolutionary governing principles that guide the development of unique metabolic signatures of organisms that share a physiological niche using a collection of 914 GENREs of pathobiont metabolism. Additionally, we show that environmental selection pressure is a possible driver of divergent evolution of metabolic function in closely related organisms, while leading to convergent evolution in distantly related organisms that occupy the same environmental niche. Further, we identify three genes informed by analysis of the GENREs that are uniquely essential to isolates of one physiological niche, the stomach. Finally, we use these three uniquely essential genes to identify and validate possible stomach pathobiont-targeted antimicrobial compounds.

### Models of pathobiont metabolism

To sufficiently capture the variation of functional metabolic phenotypes across bacterial pathobionts, we generated a collection of 914 *in silico* network reconstructions of bacterial pathobiont metabolism through an automated pipeline (Figure S1), covering 345 distinct species across nine phyla (Figure 1a,b). The scope and depth of information represented in this collection of reconstructions is substantial: across the collection there are a combined total of >1 million reactions, genes, and metabolites (Figure 1b); with individual reconstructions containing an average of about 1,500 genes, reactions, and metabolites (Figure 1d,e,f). The models were constructed using publicly available genome sequences from the Bacterial and Viral Bioinformatics Resource Center (BV-BRC)(*12*) and paired with open-source software including Python, COBRApy (*9*), and a recently developed GENRE construction algorithm (*13*), Reconstructor. We call our collection of metabolic reconstructions PATHGENN; which is the first GENRE collection of all known human bacterial pathobionts and is among the largest publicly available collections of high-quality GENREs (*14*, *15*).

**Figure 1.**
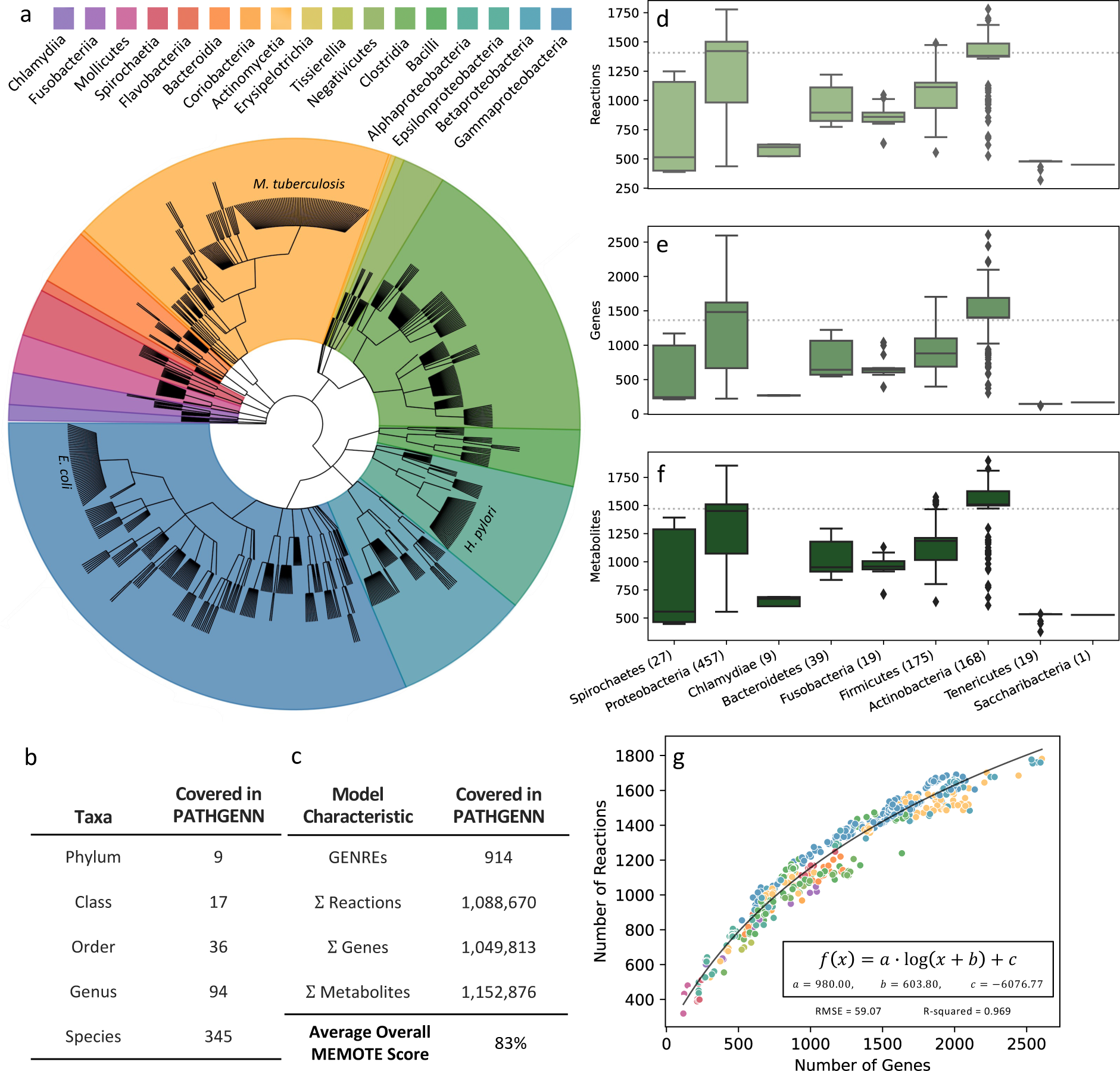
Scope of our collection of pathobiont models of metabolism. (a) Phylogenetic tree depicting the diversity of 914 considered bacterial pathobionts in the collection of GENREs. Many strains of *E. coli, H. pylori,* and *M. tuberculosis* are included in the collection, exhibits strain specificity of our collection. This cladogram was created using the GraPhlAn python tool. (b) Boxplots representing the spread of genes, reactions, and metabolites in each model, classified by phylum. The number in parentheses after the phylum name represents how many models are in that respective phylum. Dotted line in the background represents average Reaction, Gene, and Metabolite numbers across species. (c) Our collection of GENREs represents 9 phyla, 17 classes, 36 orders, 94 genera, and 345 species of pathobionts. Across the 914 models, there are a sum total of 1.09 million reactions, 1.05 million genes, and 1.15 metabolites. The average MEMOTE score across models is 83% (d) The relationship between the number of genes and the number of reactions in each model displays a positive trend and heteroscedasticity similar to other model ensembles. Colors correspond to taxonomic class of pathobiont represented by each point (same legend as Figure 1 a)

While this collection contains a wealth of metabolic and genomic information, it was important to validate the quality and relevance of this collection. To assess quality of the network reconstructions, we used the MEMOTE score, which is based on factors related to biological plausibility of the network reconstruction including stoichiometric reaction consistency, biomass reaction quality, and presence of reactions, genes, and metabolites (*16*). The average overall MEMOTE score for reconstructions in the collection is 83% (±2.5%) (Figure S2), suggesting all reconstructions in the collection are of high quality, and therefore biologically plausible.

Additionally, we determined that the relationship between the number of genes and reactions in the reconstructions is logarithmic (R^2^ = 0.97) (Figure 1g). This logarithmic trend is consistent with the expectation that there are limited evolutionary advantages for bacteria with increasingly large genomes (*17*), which further validates the relevance of our collection.

### Uncovering metabolic functions unique to a defined subset of pathobionts

To exhibit the variety of metabolic reaction subsystems present across the collection, we annotated all reactions according to the Kyoto Encyclopedia of Genes and Genomes (KEGG) and separated reactions into core (present in > 75% of GENREs), accessory (between 25% and 75%), and unique (present in < 25%) categories (Figure 2a). Most reactions across pathobionts were considered unique, which could be attributed to the large taxonomic range represented by the reconstructions. We uncovered an evident enrichment of nucleotide metabolic subsystems in core reactions, which is consistent with the ubiquitous role of nucleotide metabolism across bacterial species (*18*) (Figure 2b). Furthermore, among the unique reactions, we observed an enrichment in terpenoid/polyketide (13% more) and xenobiotic (6% more than core) metabolic subsystems. Interestingly, terpenoid/polyketide and xenobiotic reaction subsystems both relate to drug metabolism processes which can be highly variable across bacteria (*19*). Further, xenobiotic pathways are often implicated in antimicrobial resistance(*20*), suggesting that many pathogens possess unique antimicrobial resistance mechanisms.

**Figure 2.**
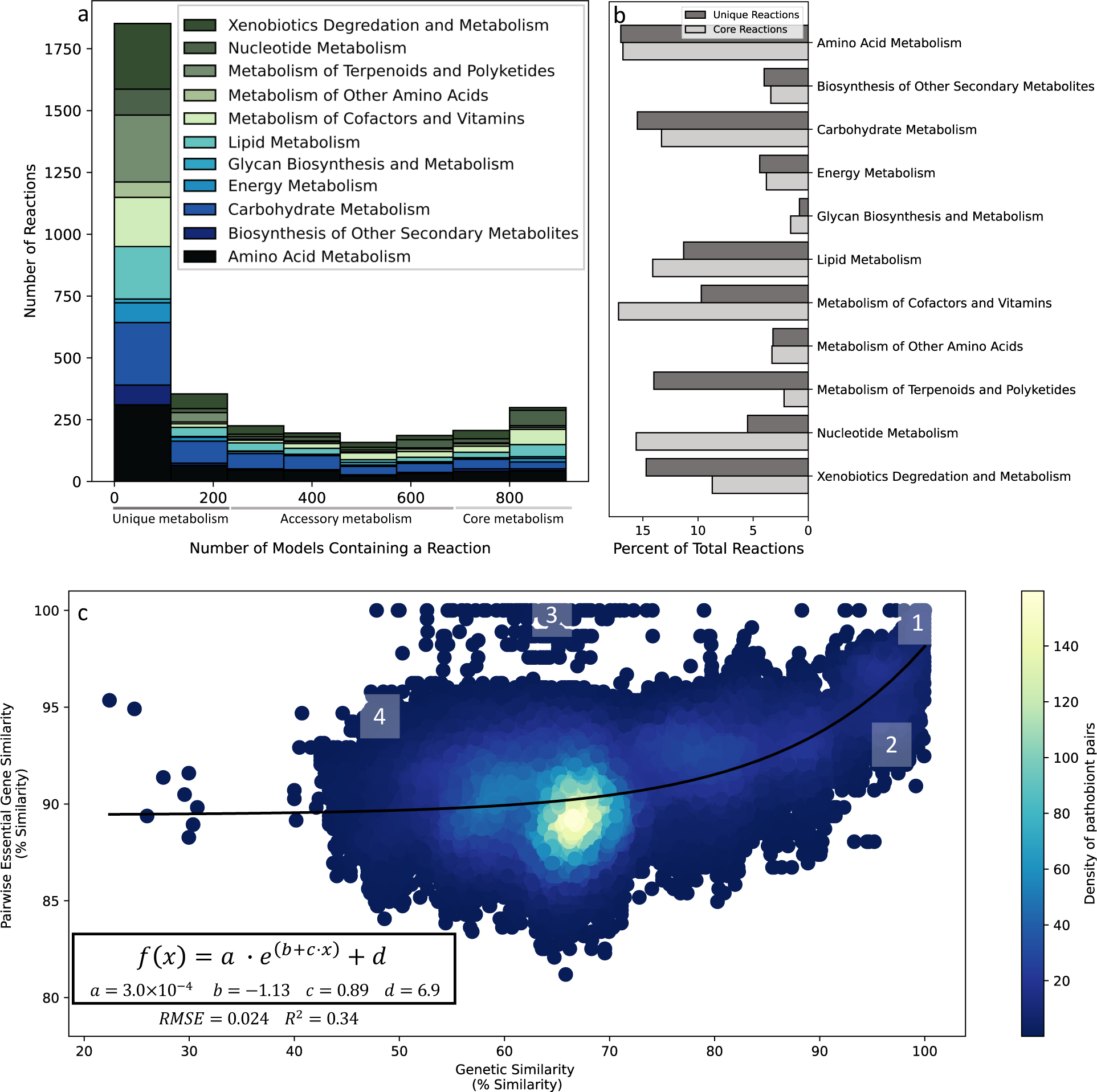
Core and unique metabolic reaction subsystems across pathobionts and the evolution of metabolic function. (a) Histogram of annotated reactions across models display prevalent reaction classes used in core metabolism (>75% models have a given reaction), accessory metabolism (between 25% and 75%), unique metabolism (<25%). (b) Enrichment of metabolic subsystems in core and unique metabolic reaction subgroups. (c) Relationship between pairwise essential gene similarity and genetic similarity in pairs of pathobionts. Trends of interest are highlighted in sections 1-4 and described in the main text. The logarithmic curve that best fits this plot is given in the bottom lefthand corner of the figure. Legend corresponds to the density of pathobiont pairs (points) in the figure.

The majority of reactions across pathobionts were considered unique to less than 25% of pathobionts showing that subgroups of pathobionts share these unique reactions, with the subgroups containing less than 229 pathobionts (25% of all pathobionts in our collection). We can think of these as metabolically unique subgroups of pathobionts, because they share unique reactions that are not present in most pathobionts. Further exploring these unique metabolic subgroups could prove beneficial, allowing us to approach antimicrobial discovery from a different perspective; leveraging shared unique functions as possible antimicrobial targets. Identifying unique metabolic subgroups and understanding the evolutionary selective pressures driving the development of these subgroups is imperative for gaining a deeper understanding of pathobiont function.

### Evolutionary drivers of unique metabolic function in pathobionts

To gain insight into the evolution of metabolism in closely and distantly related pathobionts, we analyzed the relationship of essential gene similarity and genetic similarity between pairs of pathobionts (further described in Methods). Here, we used essential gene similarity as a proxy for metabolic niche similarity, as the core essential metabolic signatures of a pathobiont provide a functional barcode for the metabolic role they play. Considering pairwise essential gene similarity as a proxy for metabolic niche, we gained insight into the dynamics of metabolic evolutionary changes across pathobionts.

We noted a strong correlation between genetic similarity and metabolic niche similarity in more closely related pathobionts (box 1 in Figure 2c). This result aligns with evolutionary expectations, similar essential metabolic functions of closely related pathobionts could be attributed to 1) their taxonomy and 2) facing the same selective pressures due to their likely physical proximity over evolutionary time. Further, it is important to consider pairs of pathobionts that are highly genetically similar, but not similar with respect to their metabolic niche (section 2 in Figure 2c).

These are pathobionts that are taxonomically very similar or even members of the same species but exhibit divergence in core metabolic functionality. A possible explanation for this divergence could be physiological niche-specific selective pressures faced by each individual member of the pair; assuming they do not inhabit the same physiological niche. For example, there has been evidence of metabolic differences in clinical isolates of *Pseudomonas aeruginosa* based on the site of isolation in the patient (*21*), suggesting environment could be a driver of unique metabolic function even within members of the same species. So, it is possible that the pairs of pathobionts shown in section 2 of Figure 2c are genetically similar organisms influenced by their physiological location to acquire evolutionarily advantageous metabolic function. Alternatively, there are situations where each member of the pair exhibits divergence in metabolic niche similarity while inhabiting the same physiological locations. In situations when closely related pathobionts are competing for resources in a shared environment, the evolution of unique metabolic function could be a strategic approach to successfully evade competition.

While there was a strong correlation between metabolic niche similarity and genetic similarity in closely related pathobionts, we saw no correlation with more distantly related pathobionts; however, there are still important evolutionary insights here. Groups of pathobiont pairs that are genetically distant but occupy similar metabolic niches (sections 3 and 4 in Figure 2c) could either be suggestive of competition or convergent evolution, depending on the circumstance. For more context, two pathobionts could be genetically distant with similar metabolic niches and inhabit the same physiological location in the host. This scenario could lead to competition since the two species are fulfilling similar metabolic niches. Additionally, due to the environmental nutrient availability and selective pressure to occupy distinct metabolic roles, pathobionts could continue to evolve and diversify through the process of divergent evolution, possibly leading to pathobiont speciation. On the other hand, there could be two genetically distant pathobionts with similar metabolic niches inhabiting completely different physiological locations in the host; this result would be indicative of convergent evolution since the similarity in essential metabolic function presumably developed independently.

These results underscore the potential of environment to shape unique metabolic function, and more importantly, show that the metabolic network reconstructions in our collection can capture complex evolutionary dynamics through constraint-based analysis. While we identified possible theoretical explanations for evolutionary dynamics in the above analysis, it was necessary to perform further analyses to support these theoretical claims.

### Functional metabolic phenotypes of pathobionts are influenced by host physiology

To provide support of these claims about the evolutionary dynamics of pathobionts, we generated and analyzed *in silico* metabolic phenotypes. We can analyze metabolic functionality through the generation of flux distributions feasible in a network reconstruction. Metabolic phenotypes are subject to evolutionary pressures like natural selection, which could result in divergent or convergent evolution and speciation. Previous studies have uncovered a strong relationship between *in silico* metabolic phenotype and evolutionary history, specifically in terms of taxonomic class (*15*, *22*, *23*). However, earlier investigations have not uncoupled metabolic phenotype from evolutionary history to consider other factors that might play a part in differentiating metabolic phenotypes. Here, we investigate the influence of pathobiont environment on the development of unique metabolic phenotypes, since our previous analysis supported a strong theoretical argument of environment being a major driver of metabolic niche differentiation.

We generated *in silico* metabolic phenotypes of all network reconstructions by utilizing flux balance analysis (FBA) to generate feasible metabolic flux distributions. To visualize individual metabolic phenotype states in the context of taxonomic class (Figure 3a) and physiological environment (Figure 3b), we used t-distributed stochastic neighbor embedding (t-SNE) (further explained in Methods). We observed significant clustering in both Figure 3a and 3b suggesting metabolic phenotype is a function of both evolutionary history and physiological environment.

**Figure 3.**
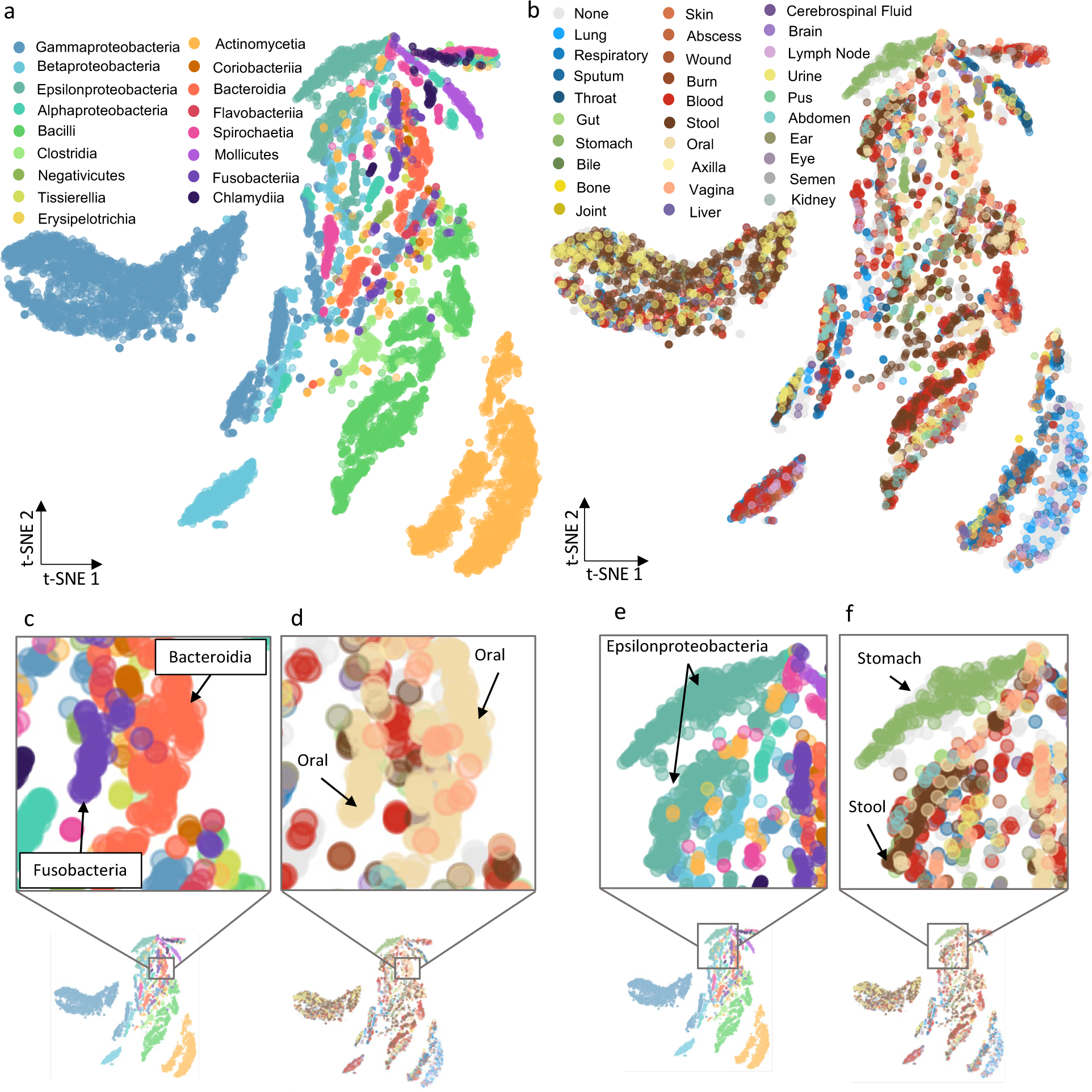
tSNE of Flux Samples Clustering on Taxonomic Class and Isolation Site. 10 flux samples across all 914 GENREs were plotted using t-SNE, and points were colored on taxonomic class (a) and physiological location(b). Areas of interest from (a) and (b) are highlighted if (c) – (f) and described in the main text.

We observed several clusters of interest in these plots, which we considered more deeply for theoretical analysis. Specifically, we noticed close local clustering of Fusobacteria and Bacteroidia species (Figure 3c). Fusobacteria and Bacteroidia are not genetically similar organisms: a multiple sequence alignment of available 16S rRNA sequences revealed significant evolutionary differences between the groups (Figure S4). Interestingly, despite having a different taxonomic lineage, these Fusobacteria and Bacteroidia pathobionts are all oral-associated pathobionts(Figure 3d) suggesting shared metabolic functionality between the groups, as evidenced by the observed clustering. Distantly related pathobionts with shared metabolic phenotypes inhabiting the same environment suggests that unique environmental properties can drive convergent evolution in distantly related organisms. This result further confirmed the observed relationship presented in sections 3 and 4 of Figure 2c, that convergent evolution is a possible explanation of genetically distant organisms with a shared metabolic niche.

Secondly, we noticed two distinct clusters of Epsilonproteobacteria, more clearly displayed in Figure 3e. The separate clusters suggest that there are two distinct metabolic phenotypes exhibited by pathobionts of this class. Interestingly, one cluster of Epsilonproteobacteria consisted solely of stomach pathobionts (Figure 3f); suggesting that these pathobionts have some distinct, unique metabolic functionality compared to their (mostly) stool Epsilonproteobacteria counterparts. This observed cluster pattern supports the observed relationship presented in section 2 of Figure 2c, that unique physiological environments can be a major driver of unique metabolic function. Biologically, this result could be explained by the unique physiological environment of the stomach: the high acidity (pH 1.5 to 2.0) (*24*) allows for only a select few bacterial species to take up residence. Further, it has been shown that *H. pylori* (a stomach pathobiont) has adapted to this extremely unique environment by utilizing metabolic pathways that taxonomically similar organisms do not utilize. (*25*).

The results from our t-SNE cluster analysis support the idea that unique physiological environment could be a driver of divergent evolution in closely related species, while driving convergent evolution in distantly related pathobionts. The divergent evolution pattern we observed is of particular interest implying that there is a selection pressure for beneficial genes that are uniquely essential to pathobionts in a specific environment. It could be possible to exploit uniquely essential genes of diverse pathobionts within a given environment to identify a physiological site-specific targeted antimicrobial therapy and thus approach antimicrobial discovery and repurposing from a different perspective; targeting uniquely essential genes that are conserved across taxonomically-distinct pathobionts that inhabit a specific physiological niche.

### Uniquely-essential genes inform stomach pathobiont-specific antimicrobial targets

Here, we leveraged the idea that environment is a major driver for the selection of unique metabolic function to exploit uniquely essential genes of stomach-associated pathobionts as potential targets for antimicrobial therapies. Targeting antimicrobial therapies to the site of infection, like the stomach, could potentially ameliorate the harmful effects of long courses of broad-spectrum antibiotics(*26*). Additionally, targeting to the infection site could reduce the need for bacterial community characterization since all organisms in the environment would be targeted. To identify uniquely essential genes to stomach-associated pathobionts *in silico*, we first determined essential genes for all strains in the GENRE collection using an FBA single-gene-knockout method in COBRApy. Subsequently, we instantiated a universal essentiality threshold to determine uniquely essential genes across stomach-associated pathobionts (see Methods).

We identified three genes as uniquely essential to stomach pathobionts (and which were not considered universally essential in any other physiological environment), *fabF*, *fabZ*, and *thyX* (Table 1). *fabF and fabZ* encode acyl-carrier-protein dehydratase and acyl-carrier-protein synthase respectively, which are both proteins belonging to the fatty acid biosynthesis II pathway. Importantly, the fatty acid synthesis II pathway is exclusively used by plants and bacteria, while animals use the type I associated pathway (*27*). This distinction makes *fabF* and *fabZ* ideal gene targets for an antimicrobial therapy since they are not involved in human fatty acid biosynthesis. Further, the fatty acid synthesis II pathway has been studied as a target for novel antimicrobial compounds due to it being ubiquitous across bacteria (*28*) . A previous study exhibited downregulation of *fabZ* expression in *Staphylococcus epidermidis* in the presence of an *α*-mangostin inhibitor, and subsequent bacterial growth inhibition was demonstrated. Further, they reported that the bactericidal action of the *α*-mangostin inhibitor is comparable with cell membrane lytic cationic antimicrobial peptides (CAMPs), suggesting this compound is a very effective antimicrobial for this application(*29*). Additionally, several studies have showed the effectiveness of cerulenin as an inhibitor of *fabF*, and have even characterized the mechanism of inhibition (*30*–*32*). In the first study citing cerulenin as a *fabF* inhibitor, cerulenin was shown to be a weak inhibitor of *Escherichia coli fabF* and showed weak growth inhibition, while being a strong inhibitor of *S. aureus fabF* with associated strong growth inhibition (*30*, *33*).

**Table 1.**
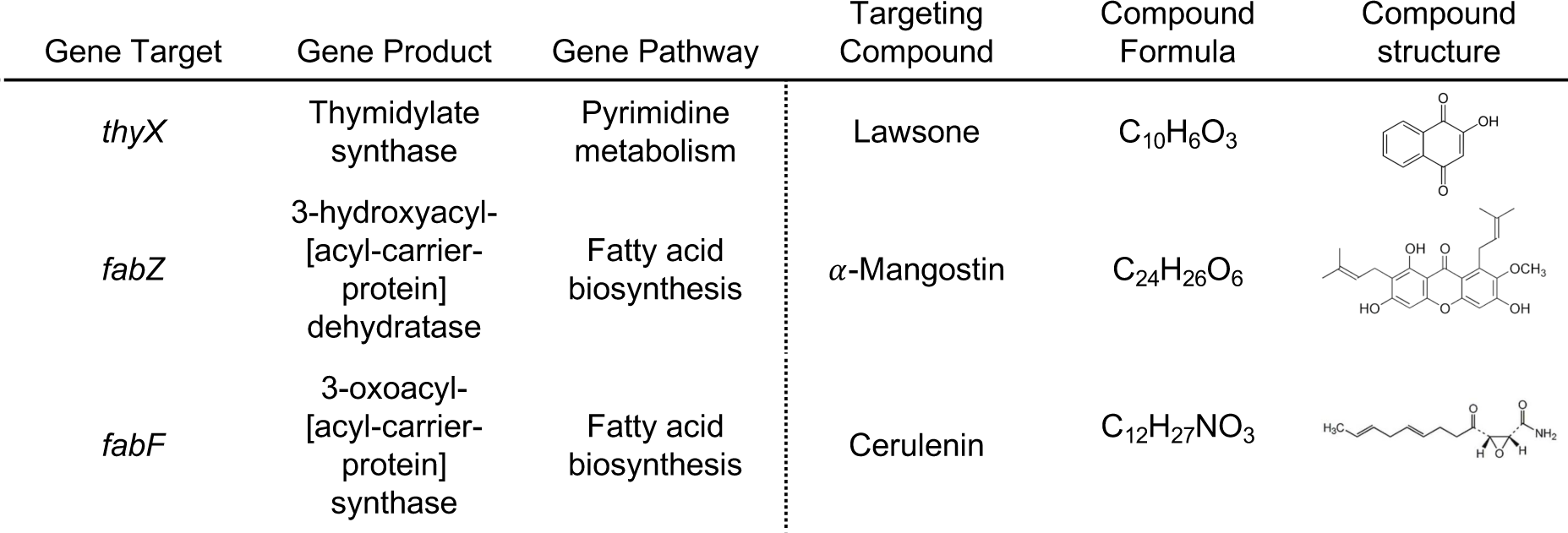
Uniquely essential genes to stomach-associated pathobionts and their corresponding chemical inhibitors. We identified three uniuqley essential genes to stomach associated pathobionts (*thyX, fabZ, and fabF)* and identified three previously verified chemical inhibitors of the genes Lawsone, *α*-Mangostin, and cerulenin respectively. Gene products, pathways that the gene is involved in, along with the inhibitor’s chemical formula, and structure are provided.

| Gene Target | Gene Product | Gene Pathway | Targeting Compound | Compound Formula | Compound structure |
| --- | --- | --- | --- | --- | --- |
| *thyX* | Thymidylate synthase | Pyrimidine metabolism | Lawson | C<sub>10</sub>H<sub>6</sub>O<sub>3</sub> |  |
| *fabZ* | 3-hydroxyacyl-[acyl-carrier-protein] dehydratase | Fatty acid biosynthesis | α-Mangostin | C<sub>24</sub>H<sub>26</sub>O<sub>6</sub> | |
| *fabF* | 3-oxoacyl-[acyl-carrier-protein] synthase | Fatty acid biosynthesis | Cerulenin | C<sub>12</sub>H<sub>27</sub>NO<sub>3</sub> |  |

The third uniquely essential gene to stomach-associated pathobionts is *thyX* which belongs to the pyrimidine metabolism pathway, specifically coding for thymidylate synthase. Thymidylate synthase is utilized in both bacterial and human cells, and is responsible for catalyzing the DNA building block thymidylate. However, the flavin-dependent thymidylate synthase, *thyX*, is only present in bacteria, and completely absent in humans (*34*), making it an optimal target for an antimicrobial therapy. There has been evidence of 1,4-napthoquinone derivatives as effective *thyX* inhibitors in *H. pylori* and *M. tuberculosis* (*34–36*), effectively inhibiting bacterial growth. Further, there has been computational evidence that many versions of 1,4-napthoquinones can inhibit *thyX,* including 2-hydroxy-1,4-napthoquinone, otherwise known as lawsone (*34*).

Our literature review corroborated predictions that *fabF*, *fabZ*, and *thyX* are essential genes for bacterial function, and provided us with possible small molecule inhibitors of each gene product. While our review did not verify that these genes are uniquely essential to stomach pathobionts, it still provided essential insights and promise that there is potential for one or all three of these genes to be uniquely essential and targetable *in vitro*. It is important to recognize that the identified inhibitors of *fabZ, fabF* and *thyX* (*α*-mangostin, cerulenin and lawsone respectively), have been shown to be effective against non-stomach-associated pathobiont species. However, this observation does not discount our hypothesis that these genes are uniquely essential to stomach pathobionts because of our definition of uniqueness (Methods). A gene can be essential to specific pathobionts in a given environment without being uniquely essential to pathobionts across the environment. For example, 1,4-napthoquinone derivatives (lawsone) were shown to be effective inhibitors of *thyX* in *M. tuberculosis*, a lung pathobiont, but that does not mean *thyX* is essential across all lung-associated pathobionts. Another point of interest is that cerulenin was reported as a weak inhibitor of *fabF* in *E. coli*, while being a strong inhibitor of *fabF* in *S. aureus,* exhibiting the variable strength of cerulenin as a *fabF* inhibitor. So, although cerulenin is not a strong inhibitor of *fabF* in *E. coli,* it is possible that cerulenin could still be a strong universal *fabF* inhibitor across stomach-associated pathobionts, following our *in silico* predictions. Therefore, it is still possible that *fabF, fabZ,* and *thyX* are uniquely essential and targetable genes of stomach-associated pathobionts. However, due to the unanswered questions that arose during our literature review and possible off-target effects of these identified inhibitors, it was necessary to validate our computational prediction further.

### Identified inhibitor determined to be stomach pathobiont-specific

To validate our computational predictions, we designed an *in vitro* assay to assess growth inhibition of stomach-associated pathobionts and non-stomach associated pathobionts subject to three known small-molecule inhibitors: cerulenin, *α*-mangostin, and lawsone. There is existing experimental evidence that cerulenin, *α*-mangostin, and lawsone are inhibitors of the genes *fabF, fabZ* and *thyX* respectively (*29–31*, *34*, *36*, *37*), which we predicted were uniquely essential genes to stomach pathobionts *in silico.* However, we wanted to validate the potential for the identified compounds to selectively inhibit growth in stomach-associated pathobionts. We selected three stomach-associated pathobionts (*Arcobacter butzleri, Helicobacter pylori*, and *Campylobacter coli* and four non-stomach associated pathobionts; *Porphyromonas gingivalis* (oral), *Pseudomonas aeruginosa* (wound), *Escherichia coli* (stool/gut), *Burkholderia cenocepacia* (cystic fibrosis lung)) as controls. We subjected each isolate to the panel of inhibitory compounds and continuously monitored growth through stationary phase (further described in Methods).

The *thyX* inhibitor, lawsone, inhibited growth in all stomach-associated isolates (*A. butzleri, H. pylori, C. coli*) (Figure 4 a-c), while not significantly inhibiting growth of non-stomach associated isolates (*P. gingivalis, P. aeruginosa, E. coli, B. cenocepacia*)(Figure 4 d-g). These results align with our computational predictions that lawsone will inhibit growth of stomach-associated isolates while not significantly affecting growth of non-stomach isolates (7/7 predictions correct (100%)).

**Figure 4.**
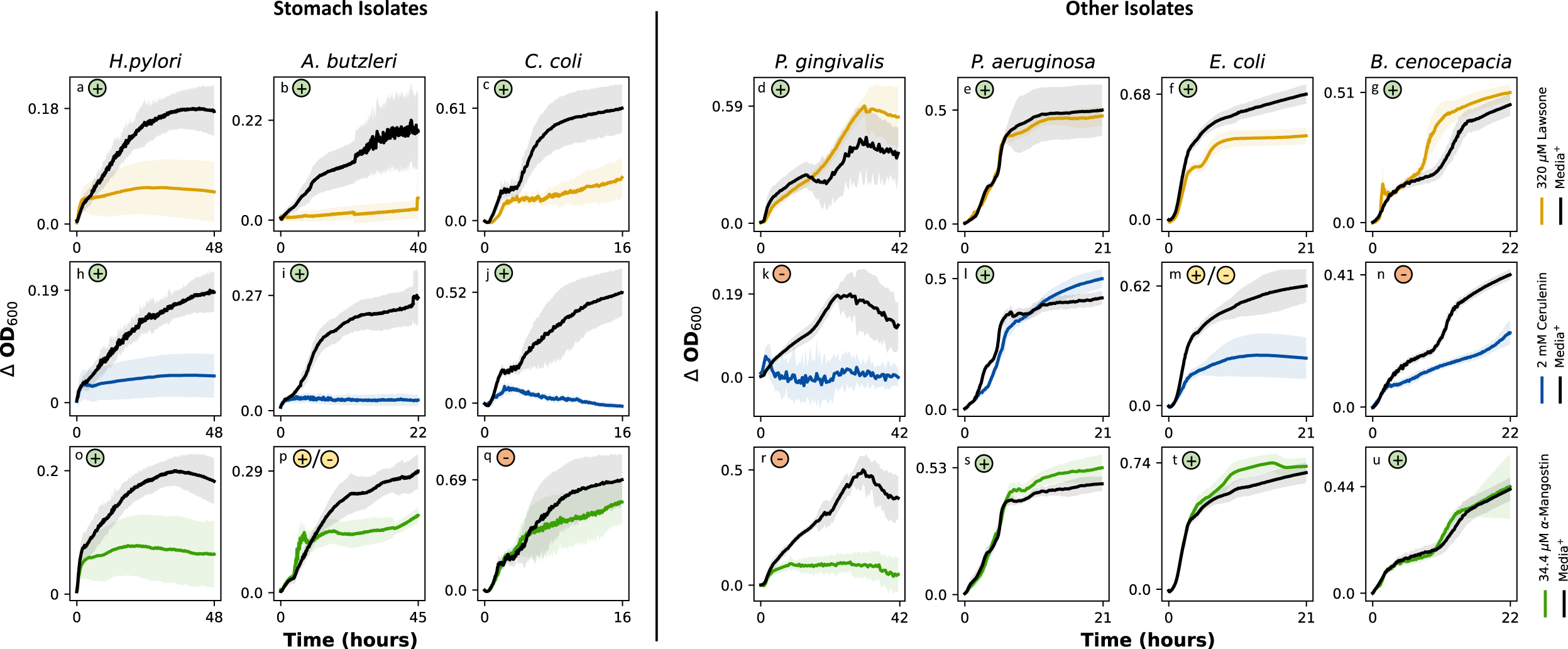
Results from validation experiments. Growth *of H. pylori, A. butzleri, C. coli, P. gingivalis, P. aeruginosa, E. coli,* and *B. cenocepecia* in the presence of three compounds; lawsone, cerulenin, and *α* – mangostin. (a-g) lawsone. (h-n) cerulenin. (o-u) *α* – mangostin.

The *fabF* inhibitor, cerulenin, inhibited growth in all three stomach-associated isolates (Figure 4 h-j), while not significantly inhibiting growth in *P. aeruginosa* (Figure 4l). However, cerulenin also showed signs of weak growth inhibition in *E. coli* (Figure 4m) and *B. cenocepacia* (Figure 4n), and strong growth inhibition in *P. gingivalis* (Figure 4k). Importantly, our result that cerulenin is a weak inhibitor of *E. coli* growth concurs with a previously published study (*30*). These results partially align with our computational predictions that cerulenin will inhibit growth of stomach associated pathobionts, but also show evidence that cerulenin is not a stomach-pathobiont-specific growth inhibitor (4/7 predictions correct (57%)).

Finally, the *fabZ* inhibitor, *α*-mangostin, inhibited growth in one of three stomach-associated pathobionts (*H. pylori*) (Figure 4o-q). Further, there was no inhibition of growth in *P. aeruginosa, E. coli, and B. cenocepacia* (Figure 4s-u). However, there was evidence of significant growth inhibition in the non-stomach-associated pathobiont *P. gingivalis* (Figure 4 r) (4/7 predictions correct (57%)). These results could be explained by the study where *α*-mangostin was cited as a *fabZ* inhibitor. *α*-mangostin was also implicated in the downregulation of many other genes, as well as upregulation of genes related to oxidative stress. So, while *α*-mangostin is an inhibitor of *fabZ*, it does not appear to be a selective inhibitor.

Interestingly, *P. aeruginosa*, *E. coli*, and *B. cenocepacia* all belong to the proteobacteria class, and all exhibit similar responses to *α*-mangostin. However, *H. pylori*, *A. butzleri,* and *C. coli* also all belong to the same class, Epsilonproteobacteria, but exhibit stark differences in responses to *α*-mangostin. This data further supports our idea that taxonomic class is not the only indicator of metabolic function, and that there are likely other factors at play driving unique metabolic functionality.

Further, our results with the *thyX* inhibitor lawsone are particularly encouraging; we inhibited growth of all three stomach-associated pathobionts while having no significant effect on growth of non-stomach associated pathobionts. With these results, we successfully demonstrated that our computational pipeline is valid for identifying unique antimicrobial targets for growth inhibition of physiological-location specific pathobionts. The idea of targeted antimicrobial therapies has been previously explored, usually applied to targeting specific species of bacteria using antimicrobial peptides (*38*, *39*). However, there has been little research on site-specific targeting of antimicrobial compounds. Developing site-specific targeted antimicrobial compounds with a data- and model-driven approach as described here could be a valuable new avenue to explore.

## Discussion

The antimicrobial resistance crisis is rapidly reducing the effectiveness of current drugs to treat microbial infections (*4*). There is a need to begin to use creative approaches to identify new or repurposed compounds that can be used as antimicrobials. Little previous work has been done to explore evolutionary pressures that lead to the rise of conserved microbial signatures, which could serve as site-specific antimicrobial targets. Here, we leveraged large-scale genomic data and metabolic network modeling to uncover evolutionary pressures driving the acquisition of unique metabolic function; which we used to identify and validate physiological niche-specific, targeted, antimicrobial compounds.

Our data-driven approach provides a valid framework for identifying highly targetable physiological locations. While our collection of metabolic reconstructions is of high quality, further curating our model simulations by providing site-specific contextualized *in silico* media conditions could enhance our analyses, potentially revealing more targetable physiological locations beyond the stomach. We believe that our computational pipeline and approach is an important beginning step for further discovery and validation of targeted, site-specific antimicrobial compounds.

## Acknowledgements

### Funding

NSF GRFP award number 1842490

the University of Virginia NIH Systems and Biomolecular Data Sciences Training Grant (grant number 1 T32 GM 145443-1

NIH R01-AI154242 and NIH R01-AT010253

National Institute of General Medical Sciences (5T32GM136615-03) NSF Grant NRT-ROL 2021791.

### Author contributions

Conceptualization: EMG, JAP Methodology: EMG, GLK, ASW

Visualization: EMG, LRD

Funding acquisition: EMG, JAP, LRD, GLK Writing – original draft: EMG

Writing – review & editing: EMG, LRD, GLK, ASW, JAP

### Data and materials availability

Additional data is available in the supplementary materials. All PATHGENN GENRE models are publicly available on GitHub along with MEMOTE benchmarking scores and all pertinent code to this study: https://github.com/emmamglass/PATHGENN.

## Supplementary materials

Material and Methods Figs. S1 to S5

## Methods

### Genome-scale metabolic network reconstruction from genome sequences

We first filtered all genome sequences in the BV-BRC (*40*) 3.6.12 database to only include those that were considered “good” quality, “complete”, and which came from “human” hosts. BV-BRC guidelines define “good” as “a genome that is sufficiently complete (80%), with sufficiently low contamination (10%)”, and amino acid sequences that are at least 87% consistent with known protein sequence. “Complete” means that replicons were completely assembled. “Human” hosts mean that the bacteria were isolated from a human host prior to sequencing.

There are 538 species of bacterial pathobionts (*2*), some of which either do not have publicly available genome sequences via BV-BRC or do not have “good” and “complete” genome sequences and were isolated from a human host in BV-BRC. For all pathobionts that pass the initial “good”, “complete”, and “human” filters, there is at least one NCBI taxid for each species, with some species having multiple unique NCBI taxids. Multiple genome sequences are available in BV-BRC for each NCBI taxid, so sequences were selected based on the presence of metadata in a hierarchical nature. This metadata requirement was instantiated because we wanted to select sequences with enough available metadata for downstream analyses. Sequences with the most associated metadata were prioritized. If multiple sequences had the same amount of metadata, we selected the sequence that had isolate environment-associated metadata. If multiple sequences fulfilled the previous requirements, the strain that had host health-associated metadata was selected. This hierarchical selection was continued for metadata categories of isolation country, collection date, and host age, in that order of priority. The resulting list contained 914 unique genome sequences. This procedure was automated with a python script available at https://github.com/emmamglass/PATHGENN.

All amino acid sequences were then automatically annotated with RAST 2.0 (*41*, *42*), and GENREs were created for each strain using the Reconstructor (*43*) algorithm. All models are publicly available (see Data Availability section). We benchmarked all GENREs using the community standard, MEMOTE (*16*), and have included overall MEMOTE scores and subcategory scores are reported in the supplementary data. Further, average overall MEMOTE scores and average major sub-category scores are shown in S2.

### Identifying core, accessory, and unique reactions and their corresponding metabolic subsystems

To identify core, accessory, and unique reactions across pathobionts, we generated a reaction presence matrix. Rows corresponded to each individual GENRE, while columns were KEGG (*44*) reactions. Reaction presence and absence was noted for each genre (1 = presence, 0 = absence). Then, a histogram was generated based on frequency of reaction presence. Reactions that were present in less than 25% of GENREs were categorized as unique reactions, reactions present in 25% to 75% or GENREs were categorized as accessory reactions, and reactions present in greater than 75% or GENREs were categorized as core reactions. Subsequently, each reaction was annotated with the corresponding KEGG metabolic subsystem to which it belongs. The histogram was then annotated with these metabolic subsystems in each bar of the histogram. Secondly, we determined the enrichment of reactions belonging to each metabolic subsystem in core reactions compared to unique reactions.

### Genetic and Essential Gene Similarity

All sequences used to create GENREs were re-annotated to determine the rRNA genome features. All 16S rRNA sequences were extracted from the annotation output, for a total of 362 16S rRNA sequences, each from a unique strain in the collection of GENREs (still representing the same 9 phyla represented in all 914 models). While there are 914 network reconstructions in our collection, only 362 of the genomes used to generate these reconstructions had annotated 16s rRNA sequences. The 16s rRNA sequences were then aligned using Clustal Omega (*45*) and the resulting Percent Identity Matrix was downloaded. Identity percentages were converted to values between 0 and 1, 0 being the most similar and 1 being the most different. Genetic distances were then converted to genetic similarities by subtracting the genetic distance from one. This value was then converted to a percentage. This metric was defined as the genetic similarity for subsequent analyses.

Essential gene profiles for each of the corresponding 362 GENREs (those with available 16s rRNA sequences) using an FBA-based, single-gene-knockout method in COBRApy(*46*) (cobra.flux_analysis.variability.find_essential_genes()). Simulations were used in a complete media context with open exchange reactions, resulting in a minimum number of essential genes. Essential genes were then converted to KEGG Orthologs, and a binary matrix was created indicating essential gene presence in each strain (1 = presence, 0 = absence). The pairwise essential gene distance was defined as the calculated hamming distances(*47*) between each strain’s essential gene profile. Pairwise essential gene distances were converted to essential gene similarities by subtracting the essential gene distance from one.

Genetic similarity vs. essential gene similarity was plotted for each pair of pathobionts. Logarithmic functions were fit to both plots using the scipy.optimize.curve_fit function in the python scipy (*48*) toolbox.

### FBA and t-SNE Dimensionality Reduction/Visualization

For each of the 914 models, Flux Balance Analysis (FBA) was performed using the COBRApy toolbox for each model in our collection to capture metabolic flux through all model reactions. 10 flux samples were taken per model for a total of 9,140 flux samples, since reducing the dimensionality of a larger number of flux samples was infeasible.

t-distributed stochastic neighbor embedding (t-SNE) (*49*) was used for dimensionality reduction and subsequent visualization of the FBA output. The perplexity parameter was selected to attempt to preserve local and global relationships in the data as best as possible, by using the relationship *P* = *N*½, where P = perplexity, and N = number of points. Points were colored based on taxonomic class, and subsequently colored on physiological location for visualization purposes.

To ensure that 10 flux samples was sufficient to capture the flux solution space as well as 100 flux samples per model would, we ran several trials of a paired-down t-SNE analyses. We randomly sampled 100 GENREs from the 914 total GENREs. Then, for each of those 100 GENREs we used 100 flux samples to perform dimensionality reduction and subsequent visualization via t-SNE (Figure S3). We performed this analysis four times, to ensure that the results would hold true for multiple randomly selected subsets of GENREs.

Through this subsequent t-SNE analysis, we still see clustering by taxonomic class in Figure S3. Specifically, we still observe large clusters of *Gammaproteobacteria* and *Actinomycetia*. Additionally, we still observe the separation of *Epsilonproteobacteria* into distinct clusters, one of which is completely comprised of stomach isolates, suggesting our larger t-SNE analysis with 10 flux samples for each of 914 GENREs is valid.

### Determining uniquely essential genes

Essential genes for all 914 models were determined using an FBA-based single-gene-knockout method in COBRApy (cobra.flux_analysis.variability.find_essential_genes()). All essential genes were translated to KEGG orthologs. Strains and their corresponding essential genes were grouped by isolation site. Essential genes present in >= 80% of strains in a given isolation source were defined as uniquely essential to that isolation source. Uniquely essential genes present in stomach isolates that are not considered uniquely essential to other isolation sites were selected, which were *fabF*, *fabZ*, and *thyX*.

### *In vitro* growth assay for computational prediction validation

We validated our computational prediction that *fabF, fabZ,* and *thyX* are genes that are uniquely essential to stomach isolates and can be targeted with cerulenin, *α*-mangostin, and lawsone, respectively. We selected three stomach isolates that were included in our network reconstruction collection, *Arcobacter butzleri* (DSM 8739)*, Helicobacter pylori* (DSM 21031) and *Campylobacter coli* (JV20). We selected four non-stomach associated isolates *Escherichia coli* (JM101)*, Pseudomonas aeruginosa* (PAO1)*, Porphyromonas gingivalis* (DSM 20709), and *Burkholderia cenocepacia* (K-56-2). Each of the isolates used is Gram negative to remove Gram status as a confounding variable.

We grew overnight cultures of each species prior to beginning each experiment. Each species was grown in a complete media to be consistent with the way computational predictions were done. However, different complete media were used for each species to optimize their growth capabilities. *A. butzleri, B. cenocepecia,* and *C. coli* were grown in Difco brain heart infusion broth (Becton, Dickinson & Co) supplemented with 5% FBS (gibco by Thermofisher Scientific). *H. pylori* was grown in brucella media (Remel) supplemented with 5% FBS. *E. coli* and *P. aeruginosa* were grown in luria broth (Sigma). *P. gingivalis* was grown anaerobically in reinforced clostridial media (ATCC medium 2107). *A. butzleri, H. pylori,* and *C. coli* are microaerophilic species, so they were grown in an airtight container with a Mitsubishi Anaeropak to keep the Oxygen concentration between 6-12% and carbon dioxide between 5-8%. All species were grown at 37° C with the exception of *A. butlzeri*, which was grown at 28° C.

Initial strong solutions of cerulenin (Sigma), *α*-mangostin (MedChemExpress), and lawsone (2- Hydroxy-1,4-napthoquinione, Sigma) inhibitors were created by first using dimethyl sulfoxide (Sigma) to solubilize each compound. Brain heart infusion media was used for subsequent dilutions to achieve the necessary final concentration.

First, we performed a minimum inhibitory concentration (MIC) assay to determine the MIC of each compound for *A. butzleri*, one of the stomach isolates. Beginning with a high concentration of each compound, we plated 2x serial dilutions of each compound with *A. butzleri* from an overnight culture. We then ran a continuous growth curve using the Cerillo Stratus plate reader, encased in an air-tight container with a Mitsubishi anaeropak to achieve the microaerophilic conditions necessary for *A. butzleri* growth. Results of the MIC assay for each compound are shown in S5. The resulting MIC for cerulenin, *α*-mangostin, and lawsone are 2mM, 34.3uM, and 320uM respectively.

The resulting MIC of each compound for *A. butzleri* was used in our final validation assay. For this assay, we subjected each species to the same concentration of inhibitors 2mM, 34.3µM, and 320µM for cerulenin, *α*-mangostin, and lawsone respectively in a 96-well plate. Each condition had eight wells containing the compound and bacteria, eight wells with the bacteria and media, and eight blank wells containing media. One plate was used per bacterial species to ensure no contamination occurred. After inoculating the plate, the plate was sealed with a Breathe-easy film to ensure gas exchange. The plate was then placed into the Cerillo stratus plate reader and monitored through stationary phase. Growth curve data was then downloaded from the plate reader via the Cerillo Canopy and saved on a local machine for analysis.

---

**S1.**
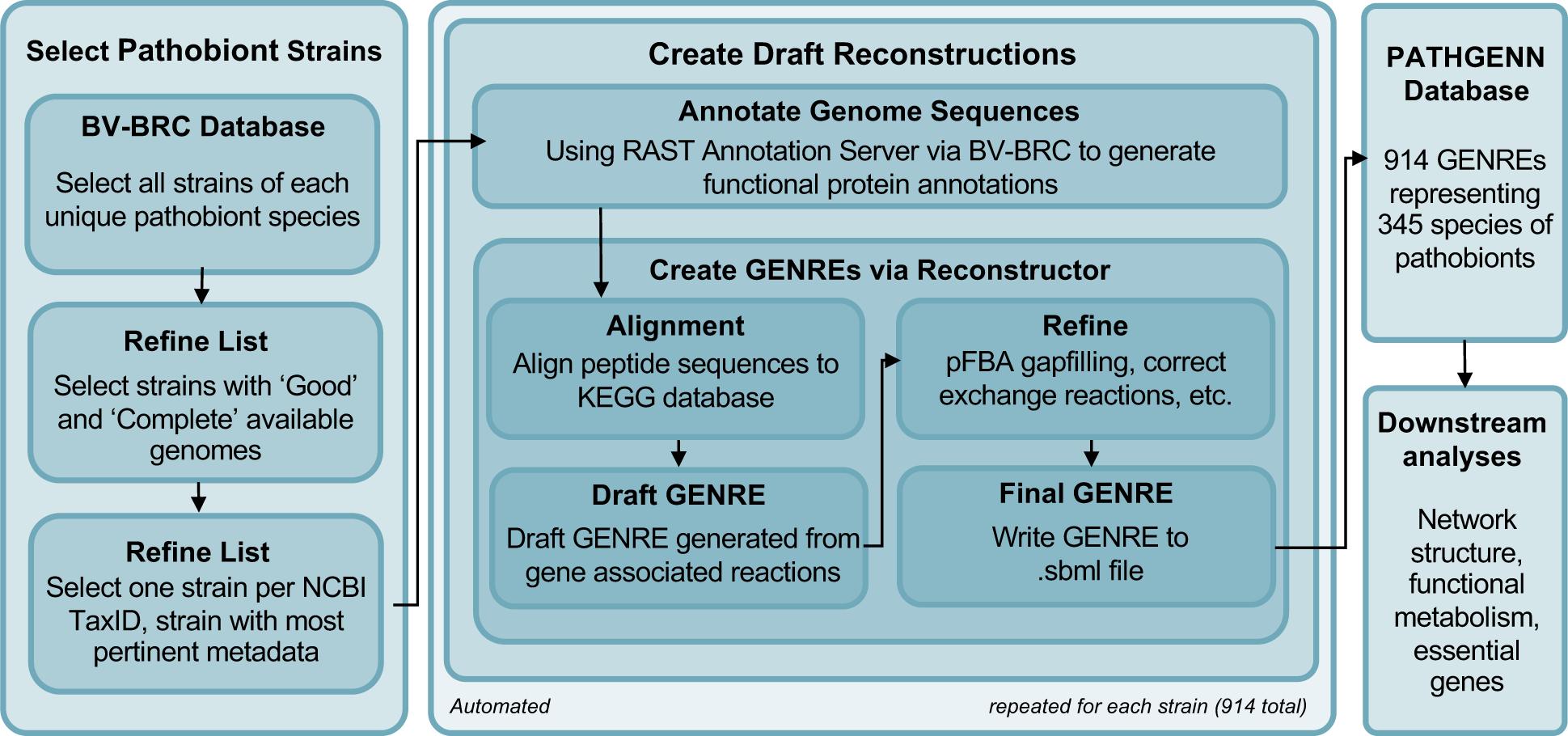
Development of the PATHGENN GENRE collection. The BV-BRC database was used to select pathobiont genome strains that satisfied quality criteria. These genome strains were then annotated using the RAST annotation toolbox to generate the amino acid FASTA file that was then used in Reconstructor to generate the 914 GENREs in the collection.

**S2.**
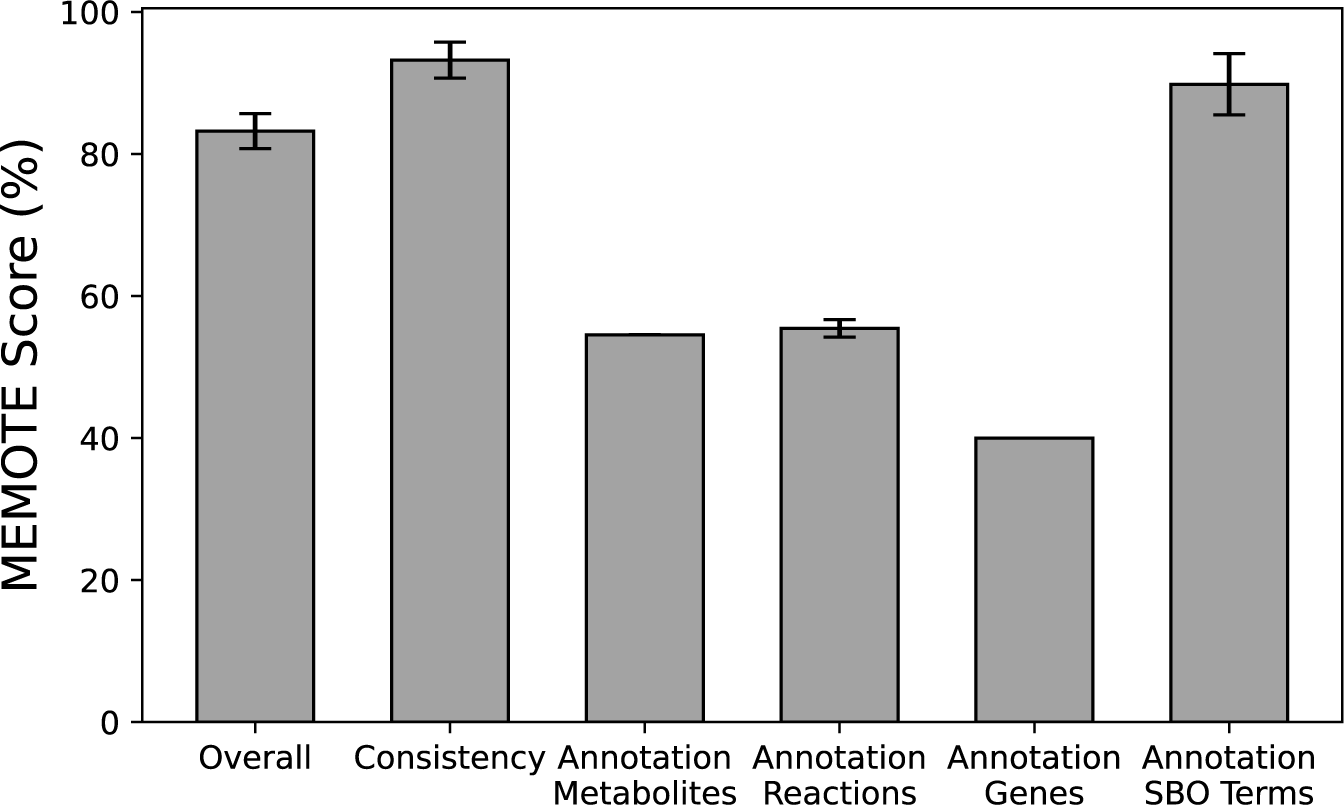
Overall and sub-category MEMOTE scores across the GENRE collection. Overall MEMOTE scores had an average of 83%, sub-category scores were also considerably high with minimal variability in quality.

**S3.**
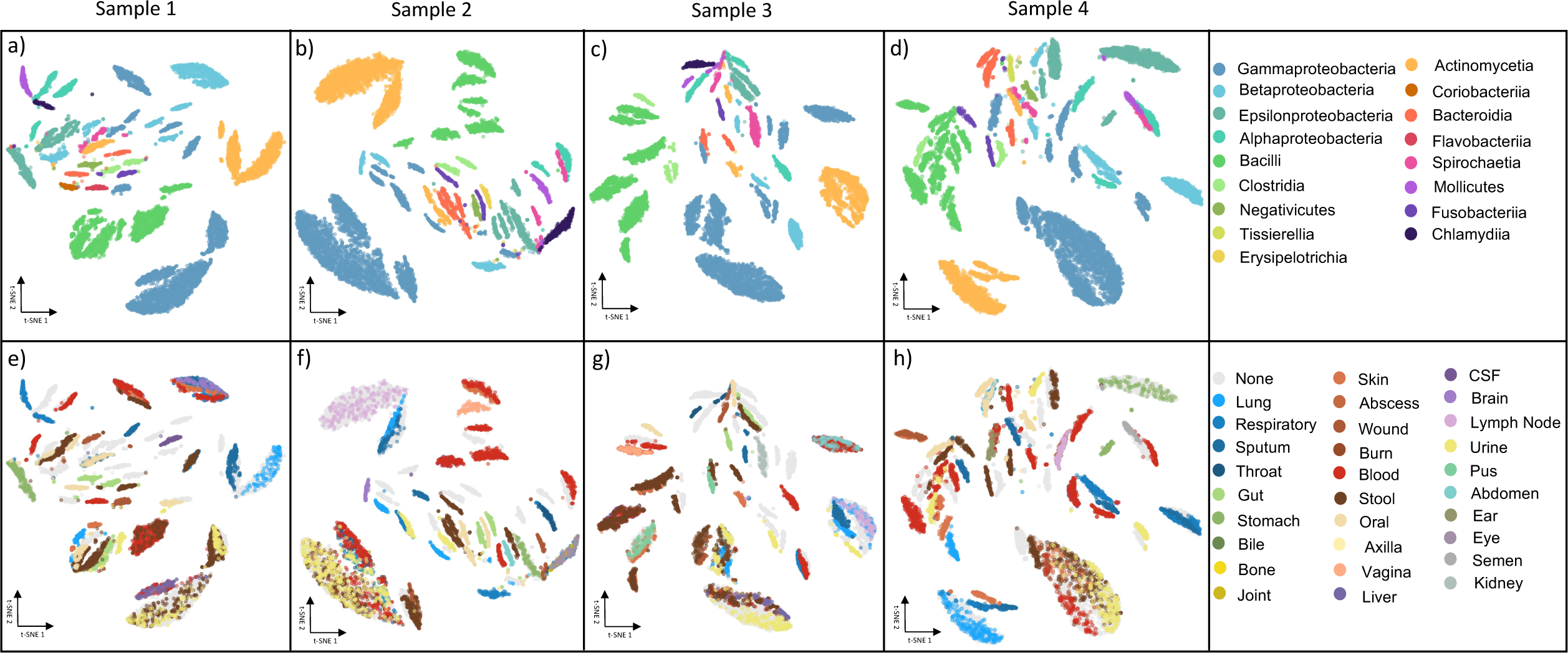
t-SNE plot of 100 flux samples for 100 GENREs. The clustering relationships seen in Figure 4 with 10 flux samples for each of 914 models are consistent with the clusters seen here with three randomly selected subsets of 100 GENREs with 100 flux samples each. Each pair of plots (a and e, b and f, c and g, d and h) represents one randomly selected subset of 100 GENREs. (a-d) are colored based on taxonomic class. (e-h) are colored based on physiological location.

**S4.**
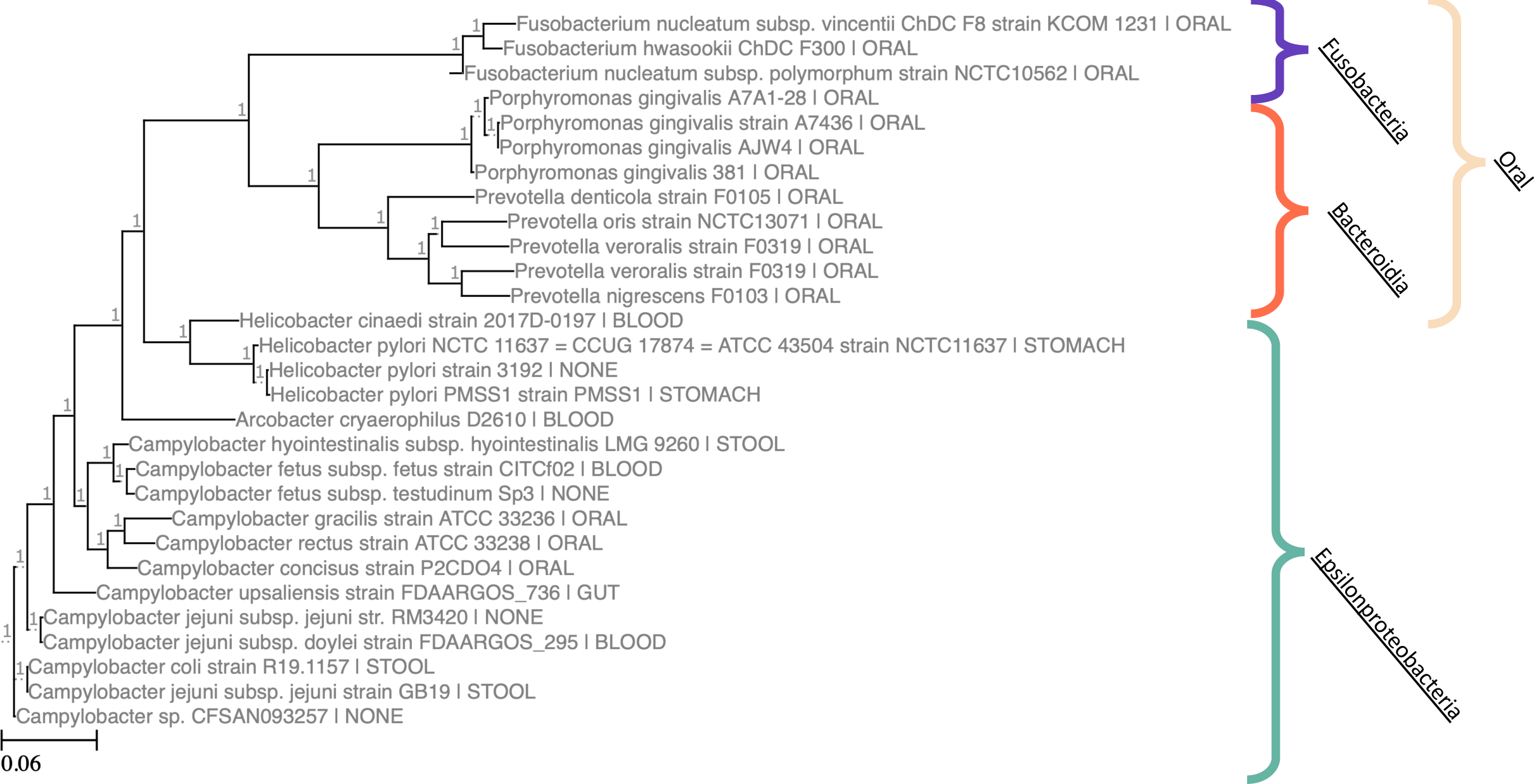
Phylogenetic tree of oral Fusobacteria and Bacteroidia species and epsilonproteobacteria species in PATHGENN with annotated 16s rRNA sequences. Fusobacteria and Bacteroidia species in the oral environment are not genetically similar. Epsilonproteobacteria are genetically similar, but occupy distinct environments.

**S5.**
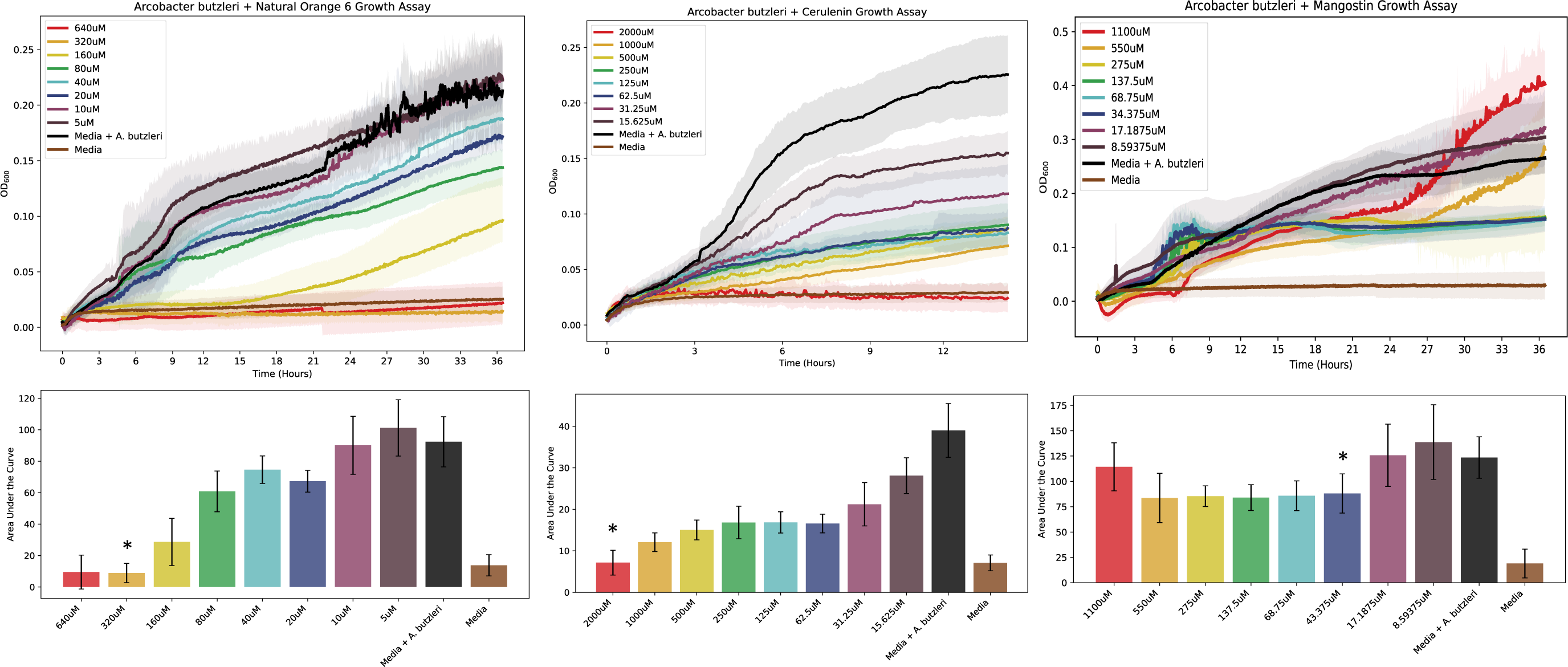
Results from MIC assay. MIC assay with *Arcobacter butzleri* for each chemical inhibitor. Stars indicate the selected MIC, the concentration used in the subsequent validation experiments.

